# GENis, an open-source multi-tier forensic DNA information system

**DOI:** 10.1101/2020.06.18.159335

**Authors:** Ariel Chernomoretz, Manuel Balparda, Laura La Grutta, Andres Calabrese, Gustavo Martinez, Maria Soledad Escobar, Gustavo Sibilla

## Abstract

GENis is an open source multi-tier information system developed to run a forensic DNA database at local, regional and national levels.^1^ It was conceived as a highly customizable system, enforcing several security policies including: data encryption, double factor identification, structure of user’s roles and permissions, system-wide secure log auditing, non-repudiation protocols and a blockchain-based option to reinforce genetic profile’s integrity. GENis is able to perform genetic profile queries of autosomal STR’s and its design follows ENFSI^2^ and ISFG^3^ standards and recommendations. In this work, we present a summary of GENis architecture, the implemented matching rule definitions, and the Bayesian framework used to provide statistical significance of cold hits, that includes a new strategy to identify common contributors in mixture pairs.

## 1. Introduction

The unprecedented statistical power of DNA technology as an identification tool has already produced a profound impact in criminal justice. In particular, the creation and development of large DNA databases have unleashed the potential of this technology to solve criminal cases [1, 2].

A forensic DNA information system stores different kind of information such as offender’s DNA profiles, genetic evidence found at crime scenes and genetic information of victims. In addition, any such system should be able to handle elimination lists of registers pertaining to personnel involved in criminal investigations, DNA-labs or chain of custody tasks. The integration of this information into a computerized environment allows implementing systematic storage and automatized comparisons among DNA profiles. This kind of systems can boost the investigation of crimes by linking DNA profiles from crime-related biological trace material to each other and/or to possible individual contributors [3- 5].

There are different IT platforms developed to run national DNA databases world-wide. In a 2016 survey, INTERPOL reported that national DNA databases were already operative in 69 member countries. The software CODIS, developed by the FBI, was reported to be used by more than half of them while other countries adopted their own developed technology to run their facilities [6, 7]. In addition, open source solutions like SmartRank have also been released to mine national databases for contributors to complex DNA profiles [8].

GENis is a DNA information system that provides data integration capabilities at regional and/or national level. It is composed by three different modules related to: (a) person identification and analysis of forensic evidence, (b) missing person identification (MPI) and (c) Disaster victim identification (DVI). In this article, we will present a detailed analysis of the functionality implemented in the first module of GENis. We will focus on system design choices, matching rules and statistical models used to assess statistical significance of detected profile matches. The paper is organized as follows: in Section 2 we summarize software design choices for the system’s architecture. In Section 3, 4 and 5 we present adopted strategies to manage the building blocks of our forensic system: marker kits, allele frequency tables and DNA profiles respectively. In particular, we explain the *group* and *category* classification system implemented in GENis to provide maximum flexibility in connection with stringency and matching rule definitions. In Section 6 we summarize the Bayesian framework used to assign statistical significance to profile associations. Simulation results were also included in order to quantify the statistical power of the system. Section 7 describes how GENis organizes matching results for active queries and introduce a scenario-testing tool designed to aid the user to weigh different hypothesis. In addition., we summarize in this section the system-wide notification circuit between involved parties to convert a hit into a match. In Section 8 some considerations about GENis multi-tier deployment are presented. Finally, discussion and conclusions are drawn in Section 9.

## 2. Software architecture

The GENis system exclusively relies on open source technology (see Table 1). The server was implemented in JVM8 and the Scala functional programming language. For security and performance reasons, information is stored in two different databases: a) A relational PostgreSQL database that stores system configuration, DNA profile metadata and the operational log database and b) A non-relational MongoDB database that stores DNA profiles and executes matching queries taking advantage of MapReduce operations.

**Table 1.**
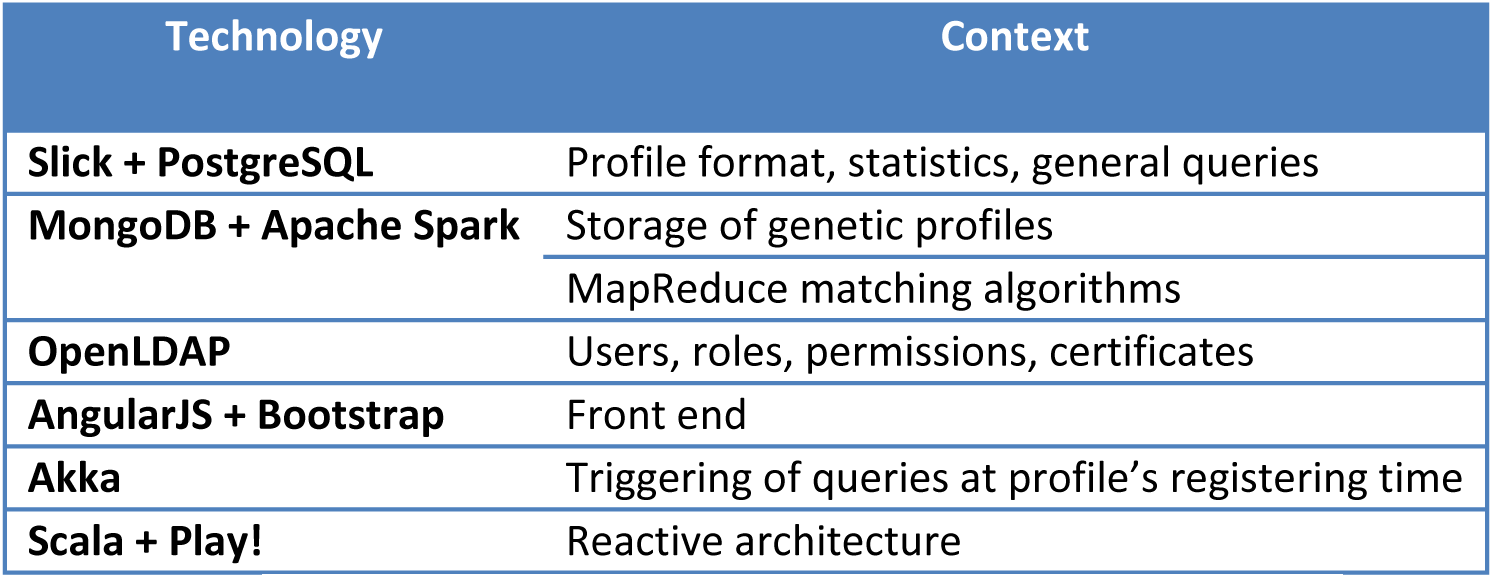
Summary of open source technologies that are used in GENis.

GENis interface was built following the single-page application pattern. In this type of applications, the browser executes the front-end component following a model–view-controller framework, and communicates with the server through remote services. The web presentation is generated using AngularJS visual components, and Bootstrap for laying them out. The front-end application communicates with the server through REST/JSON services. The server of the application was built on Play! Framework 2.3.x.

We implemented a double authentication strategy to warrant system accessibility: a username and password combination should be provided at login, along with a one-time password generated from a shared secret key and the current time. This time-based one-time password is provided by the Google Authenticator App. Moreover, user’s roles can be defined in order to restrain the operations each user can perform in the system. Table SM-T1 in supplementary material lists the kind of permissions that can be granted to different user’s roles. Further details on user registration and authentication procedures can be found in Section 2 of the User Manual (UM) included as Supplementary Material.

Additionally, an optional functionality allows checking the integrity of genetic profiles through the generation of cryptographic signatures and their inclusion in an immutable distributed ledger administered, for instance, by a national blockchain authority.^4^ That authority can guarantee the immutability of each stored record, in such a way that the authenticity of all active profiles within the database becomes verifiable. Thus, the blockchain optional functionality brings the possibility of verifying that a genetic profile has not been modified since inception in the database.

## 3. KITS and Markers

Most of the STR markers and kits that are currently used in forensic analysis laboratories can be employed in GENis. Currently, 39 autosomal kits are included (see Table SM-T2 for a summary of STR markers and kits). In addition, new STR markers can be included and custom kits can be defined upon them (see Sections UM-8 and UM-9 for further details).

## 4. Allele frequency tables

Statistical significance assessments (e.g. the estimation of random match probabilities) rely on observed allele frequencies reported for the population of interest. GENis provides CRUD capabilities to Create, Read, Update and Delete allele frequency tables that are looked up for probabilistic calculations. Whenever a population allelic frequency table is created or uploaded into the system, users should specify how genotype probabilities should be computed from observed allelic frequencies (the genotype probability model used by GENis is summarized in Section SM-1 of the Supplementary Material). For instance, either the NRC II recommendation 4.1 or the NRC II recommendation 4.10 scheme [9] should be selected, appropriate kinship coefficient values should be specified and a strategy to assign frequency values to rare new alleles should be chosen (see SM-2 for details). Section UM-10 describes implementation and operational details.

## 5. Profiles

### 5.1. DNA profile model

GENis uses a binary model to represent DNA electropherograms (see SM-3 for details). DNA profiles can be automatically imported from *GeneMapper’s* (Applied Biosystems, USA), combined-table exported text files. Alternatively, profiles can be introduced manually using a double-blind procedure. Metadata, such as court case information, type of biological sample, etc, can also be input into the system. Importantly, every entered profile should be assigned to a given profile-group and profile-category. These features lay down important aspects of the profile usage and management inside GENis, like admissibility, stringency criteria and specific matching heuristic rules (see next Section).

Quality assurance policies for DNA-profiles are necessary to promote a reliable database performance (see Sec 3.5 and 3.9 of [4]). Definitions and recommendations provided by expert committees about the admission criteria for low template DNA samples and/or mixtures of more than 2 or 3 contributors, can be accommodated and implemented in the system. Apart from the chosen criteria, GENIs follows the strategy proposed by Haned and collaborators to estimate the number of contributors of a given mixture that can be used for subsequent analysis (for the sake of completeness we summarized the followed approach in SM-5).

A given GENis instance could incorporate DNA profiles coming from different laboratories. The system provides CRUD capabilities to manage information from each contributing lab. In particular, estimations of *drop-in*, and *drop-out* parameter values, to be used in LR calculations, could be specified for each lab.

### 5.2. Profile groups and categories

Profile groups and profile categories are defined in GENis in order to implement a customizable profile organization scheme that permits to manage the system behavior in connection with admissibility criteria, association rules, and matching procedures.

The *group* classification is meant to differentiate DNA profiles coming from stains obtained at crime scenes from those profiles obtained from known sources. The former could be indexed, for instance, as belonging to the ***U****ncertain* group, *U*, whereas the second ones as belonging to the ***C****ertain* group, *C*.

Each profile should be further characterized as belonging to one of system-wide categories that can be defined to fine-tune the logical processing rules that will affect it. The categorization scheme is fully customizable and can be adapted to different working protocols and policies. Table 2 below shows a possible categorization system.

**Table 2.** Possible categorization scheme for DNA profiles. n is the estimated/assumed number of contributors.

| Group | Category | Description |
| --- | --- | --- |
| **C** | S | DNA profile of a suspect |
|  | V | DNA profile of a victim |
|  | D | DNA profile of putative cross-contaminant contributor |
| **U** | E1 | DNA profile from crime scene stains (n=1) |
|  | E | DNA profile from crime scene stains (n>1) |
|  | Ek | DNA profile from crime scene stains (n>1), with known contributors |

Different admissibility requirements can be independently set for each profile category regarding, for instance: the minimal number of informative loci or the maximal number of loci showing trisomy. In addition, profile association rules can also be defined, for instance, to link *V*-type profiles with *Ek* profiles for situations where a profiled victim is one of the known contributors to a crime scene mixture evidence.

Matching rules can be defined to customize the queries that are automatically triggered whenever a new profile of given category is incorporated into the system. It is possible to select several target categories against which the new profile should be compared at entry/creation time. For each of them, it is possible to specify the kind of matching algorithm that should be used (see below), the minimal number of loci displaying concordant allelic values, the maximal number of non-concordant markers that will be tolerated and the stringency criterion of the search. GENis follows the three-level stringency scheme (i.e. high, moderate, low) suggested by ENFSI in Section 5.4.2 of [10] (see SM-6 for details). Every profile included in the database is in anactivestatus by default. This means that it will participate in every matching procedure affecting its own category. The removal of a given profile from the database is implemented as a logical procedure that irreversibly changes its status to an *inactivated* one. In this new state the profile is simply ignored for future queries and cannot return to its previous state. Eventually a hard deletion protocol could be incorporated in accordance to specific legal frameworks governing expungement and profile removal polices.

Further functionality and operational details regarding profile groups and categories can be found in Section UM-7.

### 5.3. Profile matching

Figure 1 depicts different kind of matching rules that could be established between different profile categories. There are 3 kinds of matching strategies to achieve the following different tasks: person identification (I), establishment of contributor status in a given evidence (C), and identification of a common contributor in pair of uncertain samples (CC). In addition, the information of the victim associated profile, *V*, for the *Ev* case is used to define obligated alleles in the search heuristics (A).

**Fig 1.**
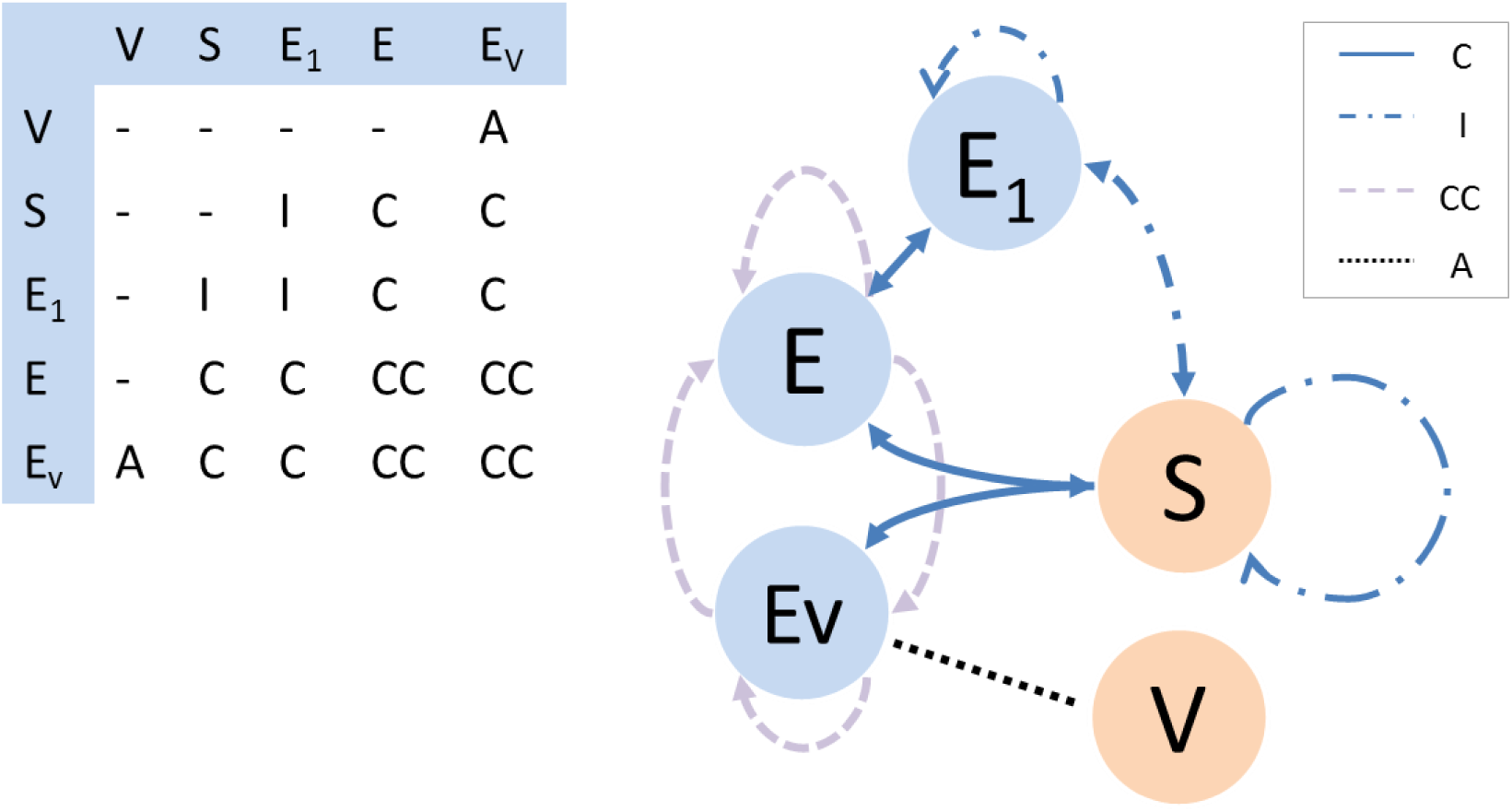
Purpose underlying match detection between profiles of different categories (described in Table 2). I: identification, C: contribution, CC: identification of a common contributor. A: association link. Right panel: Metagraph of searching/matching rules. Nodes represent profile categories. The dotted line represents an association between *Ev* and *V* categories used to identify *obligated* alleles in search heuristics. Dashed arrows represent queries that aim to identify common contributors in evidence samples (CC). Solid arrows represent matching rules with specific stringency criteria to identify contributors of a given evidence (C), and dot-dashed arrows represent matching a strategy to asses for the identification of contributors. The adjacency table of the graph is included as an inset.

When a new profile enters the system, queries are automatically triggered according to the pre-defined rules. For instance, according to the query rules depicted in Figure 1, a new suspect profile (*S* category) will be compared with already stored profiles of category *E*_*1*_, *E* and *Ev*. A match will be reported considering stringency criteria (see SM-6) defined for each kind of comparison. In this way, a “high stringency” matching level could be employed for *S*<*-*>*E*_*1*_ associations, as an identification task lies behind a match detected between profiles of such categories. On the other hand, a “moderate stringency” level could be appropriate for the matching between S and E category profiles, as a *contribution* assumption is relevant in this case.

## 6. Genotype probabilities and likelihood calculations

GENis leverages on the statistical framework developed by Curran and collaborators [11] to estimate the probability of observing DNA evidence under different hypothetical scenarios (for completeness purposes, a brief overview of the methodology is included in SM-4). The implementation followed the one considered in LRmix Studio and its R code *forensim* package, a renowned open-source software suite developed for the interpretation of forensic DNA mixtures [12-14].

### 6.1. Complex DNA profiles

In this section we present a simulation study to assess for the statistical power of GENis methodology to identify contributors to multiple-donor DNA profiles. We considered allele frequencies for 15 autosomal short tandem repeats loci for the American Caucasian population [Butler2003]. CSF1PO, FGA, TH01, TPOX, vWA, D3S1358, D5S818, D7S820, D8S1179, D13S317, D16S539, D18S51 and D21S11 markers belonged to the core CODIS loci used in the US, whereas D2S1338 and D19S433 belonged to the European core loci. We adopted a simulation strategy similar to the one used by Benschop *et al* to validate the Smartrank software [8].

Using allelic probabilities, we first simulated, with the aid of the *forensim* R package [12], 25 reference profiles denoted ‘seed genotypes’. For each one of them we generated a set of *resembling* genotypes considering almost identical copies of the original profiles, but for a single allele of a randomly chosen locus changed for a *rare allele* (we assumed *P*_*rare*_ = 2.4 10^−4^). We also generated 30 additional profiles that shared 100% (10), 75% (10) or 50% (10) of alleles with two or three-donor mixtures generated using the *seed* genotypes (*from-mix* profiles). We additionally considered, for each *seed* genotype, a parent-like and a brother-like profiles. Finally, 200 independent random genotypes were sampled from the population. Overall, 330 profiles were simulated (25 seed, 25 rare-copies, 50 familial, 30 from mixtures and 200 independent profiles).

As a gold standard we considered ten mixture profiles generated from two and three known seed profiles respectively. Drop-out altered profiles were simulated for each one of them randomly removing 0%, 20% and 50% of their allele content to model none, moderate or severe drop-out situations. Each of the 330 profiles of the simulated database were then examined under a prosecutor and defense hypothesis and corresponding LR values were estimated. The true number of contributors, a fixed drop- out rate of 0.01, a drop-in probability of 0.05 and a *θ* -correction value of 0.01 were considered in the calculations.

In Table 3 we reported the number of cases displaying LR values greater than one, as this is a first order minimal necessary condition for profile identification. For two-donor mixtures, a complete retrieval of the right seed-profiles was achieved for none and moderate simulated drop-out levels. On one hand, 20% of the sought profiles were missed for the severe drop-out situations. The retrieval of seed-profiles for 3-donor mixtures was also successful in absence of drop-outs, albeit some performance degradation was observed for moderate dropout levels. On the other hand, high dropout situations severely impeded the identification process for three-donor mixture. It can also be seen from the table that several *rare* profiles, that differed just by one allele from the queried genotype, also presented LR values larger than unity.

**Table 3.** The table reports the number of sought (S), rare (R), from-mix (M), father-like (F), brother- like (B) and population (P) profiles having LR values greater than unity. Five mixture samples, coming either from two- and three-donors, were considered for a given drop-out level. The AUC_01_ column shows the estimates of area under the ROC curve (see text). Values of this quantity were normalized and lay in the [0,1] interval. A value of unity means that every single sought seed-profile was ranked before the other type of analyzed profiles.

| Contributors | Drop-out | S | R | M | F | B | P | AUC<sub>0.1</sub> |
| --- | --- | --- | --- | --- | --- | --- | --- | --- |
| **2** | 0% | 10 (100%) | 7 (70%) | 1 | 0 | 3 | 0 | 0.995 |
|  | 20% | 10 (100%) | 6 (60%) | 1 | 1 | 1 | 0 | 0.995 |
|  | 50% | 8 (80%) | 3 (30%) | 0 | 0 | 0 | 0 | 0.949 |
| **3** | 0% | 15 (100%) | 5 (30%) | 0 | 0 | 2 | 0 | 0.994 |
|  | 20% | 11 (73%) | 2 (13%) | 0 | 0 | 2 | 0 | 0.973 |
|  | 50% | 4 (27%) | 1 (6.7%) | 0 | 0 | 0 | 1 | 0.922 |

Figure 2 displays LR estimations for the five 2-donor and the five 3-donor mixtures in the upper-left and upper-right panels respectively. Each subpanel shows data for the three dropout levels considered for each mixture-profile. Red, orange, violet, light-blue, blue and grey colored circles represent seed, rare, from-mix, father-like, brother-like and population independent profiles. Horizontal lines signal the LR=1 level. Lower left and right panel display ranking values aggregated by drop-out simulated levels for two- and three-donor mixtures respectively. It can be seen for 2-donor mixtures (upper-left panel) that the sought seed genotypes typically presented LR values larger than unity by several orders of magnitude. Some performance degradation can be observed for increasing drop-out levels. However, the right genotypes were typically over-represented in top-ranking LR positions. In order to quantify this trend, we considered results in rank space and estimated the area under the receiver-operator curve (ROC), between 0.9 and 1 specificity boundary levels (AUC_0.1_). The ROC curve serves to illustrate the ability of a binary classifier system (e.g. sought profile or not) in terms of sensitivity and specificity, parameterized by the considered classification threshold (LR value larger than a given value). The normalized partial AUC statistics shown in Table 3 is commonly used as a summary measure of the receiver operating characteristic (ROC) curve. It ranges from 0 to 1, one being a perfect classifier that ranks test cases on top of control cases [15, 16]. Despite the sought genotypes still presented the largest LR values for 3- donor samples (bottom-right panel and AUC_01_ values reported in Table 3), a general decline of absolute LR values can be observed for these cases (upper-right panel, and column S of Table 3). These findings warn against the use of poor-quality 3-donor samples for identification purposes.

**Fig 2.**
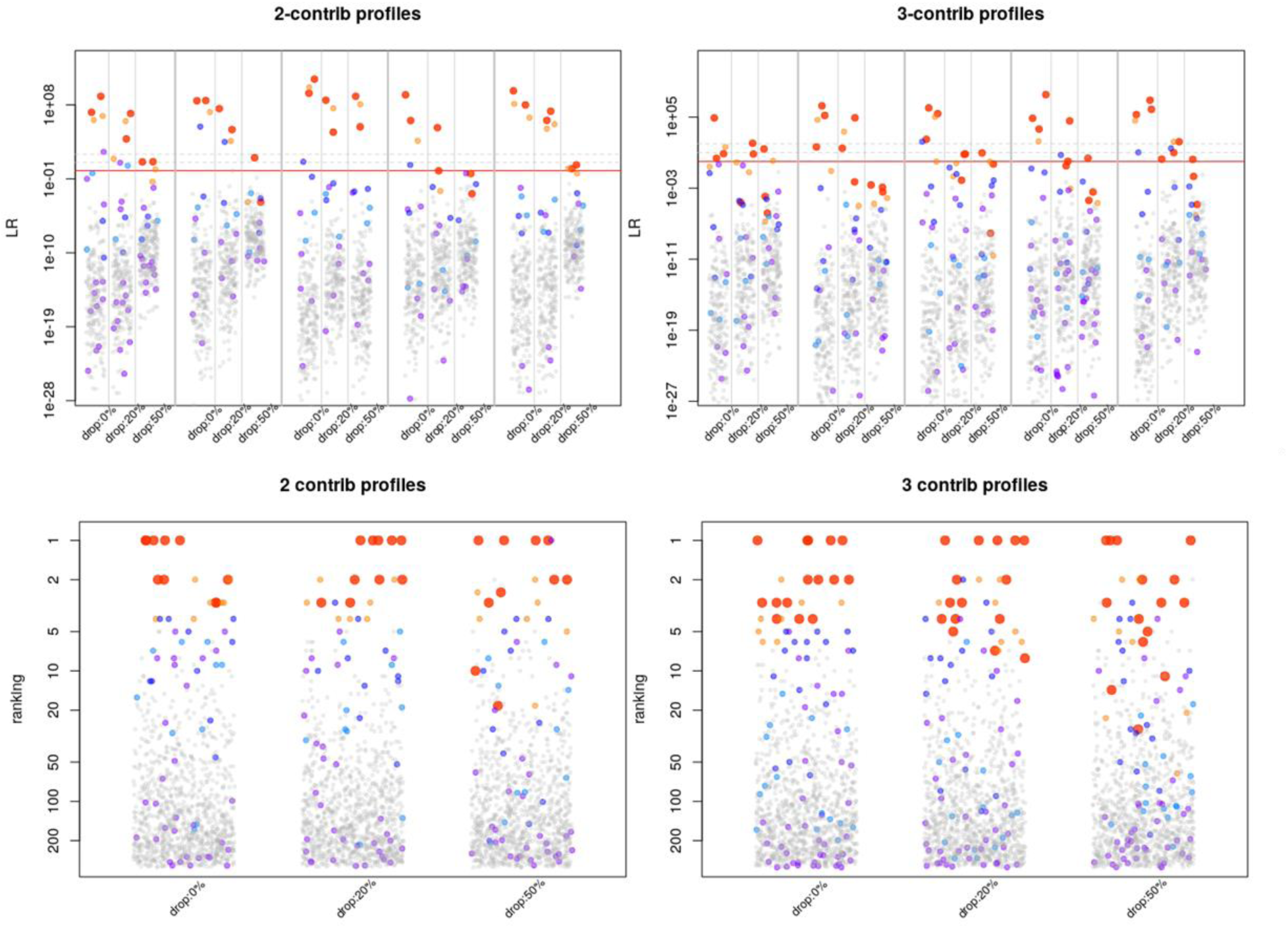
Upper-left and upper-right panels show LR estimations for the five 2-donors and the five three-donors mixtures respectively. Each subpanel shows data for the three dropout levels considered for each mixture-profile. Horizontal lines signal the LR=1 level. Lower left and right panel display ranking values aggregated by drop-out simulated levels for two- and three-donor mixtures respectively. Red, orange, violet, light-blue, blue and grey colored circles represent seed, rare, from-mix, father-like, brother-like and population independent profiles respectively.

### 6.2. Common contributor assessment

GENis implements a novel methodology to assess for the statistical significance of the identification of common contributors in DNA mixtures. Differently from recent presented procedures [17], it was specifically developed for binary profile models. Mathematical derivation of the likelihood statistics (*L*_*x*_) was included in Section SM-7.

We relied on numerical simulations in order to quantify the statistical power of the common contributor LR_x_ statistics (Equation [21] Section SM-7). For this exercise, we considered allele frequencies^5^ for 15 autosomal systems included in the Powerplex-16 System (CSF1PO, D13S317, D16S539, D18S51, D21S11, D3S1358, D5S818, D7S520, D8S1179, FGA, Penta D, Penta E, TH01, TPOX, and vWA). First, we simulated 1000 individual reference samples. We further sampled triplets of individual profiles to produce 1000 pairs of 2-person mixtures presenting one common contributor. In addition, we generated another 1000 control pairs of independent mixtures from 4 different individual profiles. We estimated the LR_x_ statistics for these samples using the formulae presented in SM-7, considering *p*_*dropin*_*=0*.*005, p*_*dropout*_*=0*.*01* and θ*=0*.*02*

In order to statistically assess whether a common contributor has participated in a given pair of mixtures, GENis considers all pair combinations (with repetitions) of shared alleles to build the set of putative common contributors. In connection with this, we noticed that in our simulations the large majority (>70%) of the generated independent control pairs did not present common alleles in at least one locus. As LR^(s)^ of different loci combine multiplicatively (see Eq 21 SM-7), null composite LR_x_ were produced for these cases. The bottom panel of Fig 3 shows (red square symbols) that the percentage of null LRs found in control pairs increases as a function of the number of considered systems, leveling off at around 12 loci. This means that finding shared alleles in two independent 2-contributor mixtures, for all 15 markers is in fact a rather non-trivial restriction.

**Fig 3.**
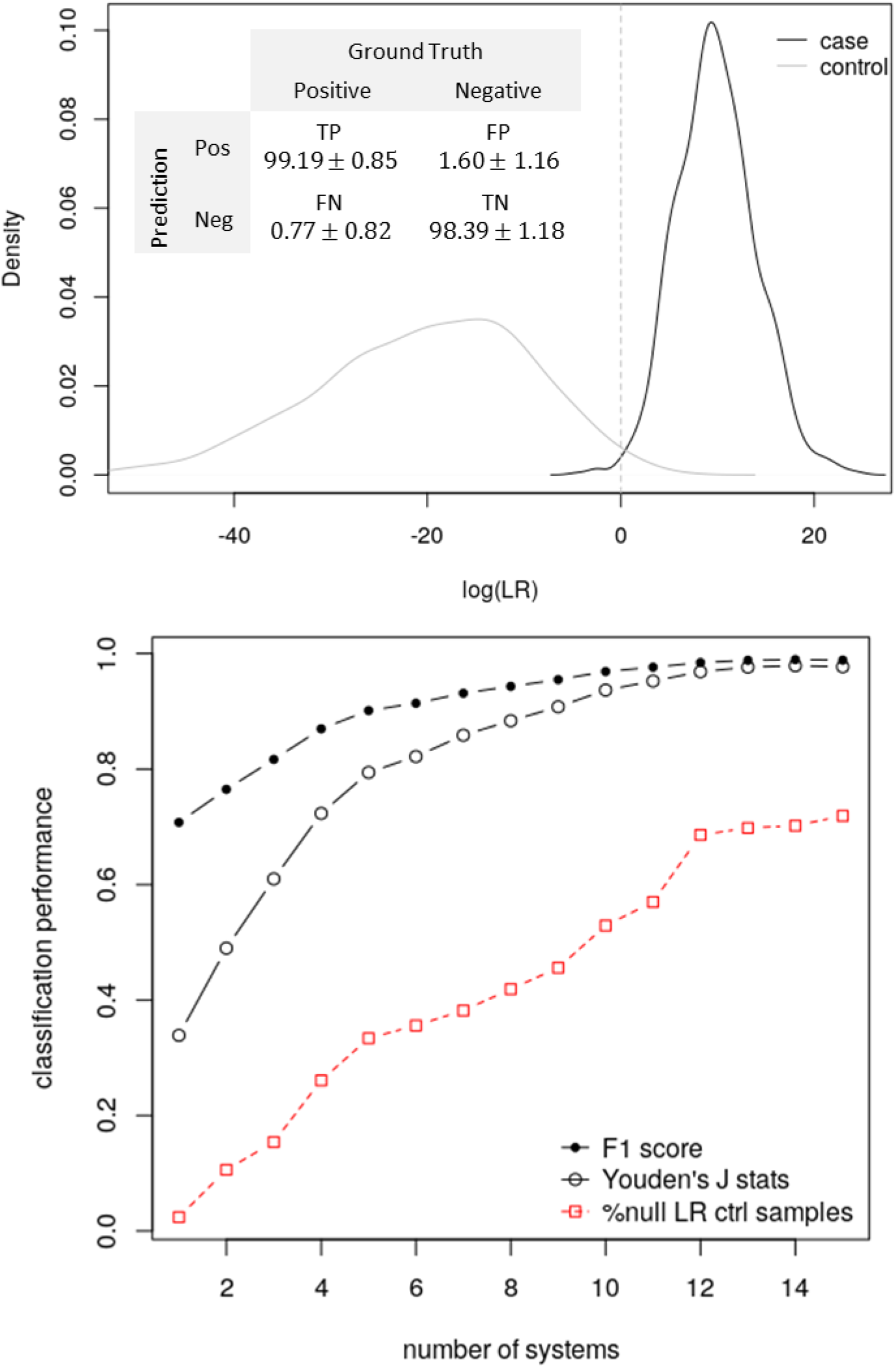
Upper panel: Density distribution functions for the *LR*_*x*_ statistics are shown for the ensemble of 2-person profile pairs having a common contributor (black) and control independent 2-contributors profile pairs (gray). The respective confusion matrix is displayed as an inset. Lower panel: Classification performance of the LR_x_ statistics to discriminate case and control profile pairs, as a function of the number of considered markers. F1 and Youden’s J statistics are shown using solid and empty black symbols respectively. The fraction of null LR_x_ obtained for independent 2-contributors profile pairs is shown using red empty squared symbols and a dashed red line.

Figure 3 upper panel displays log LR_x_ density distribution functions for truly associated 2-contributor samples (black lines) and control independent mixture pairs (gray lines). We considered only non-null LRs for control cases and, accordingly, reduced the number of systems considered for non-control common contributor cases to produce fair comparisons. It can be appreciated from the figure that control cases consistently present LR_x_<1 values, whereas the opposite is true for associated mixtures. The confusion matrix, in the figure’s inset, shows that for a threshold value of LR_x_*=1, the consolidated LR_x_ statistics for 15 systems achieves large statistical power (TP=99.12±0.85) at negligible levels of Type I and Type II errors (1.60±1.16 and 0.77±0. 82 respectively).

The classification performance of the LR_x_ statistics (LR_x_*=1) as a function of the considered number of markers is shown in the lower panel of Figure 3. The F1 score and the Youden’s J statistics (see [18] and Section SM-8) are displayed using solid symbol/continuous black lines, and empty symbols/dotted lines combinations respectively. It can be seen that near perfect scores for both figure of merits were achieved when around 12 markers were considered.

We also extended our analysis to study the ability of the association test to link 2-person and 3-person mixtures through the identification of a putative common contributor. In this case our simulations showed a 20% decrease of the statistical power of the test (TP: 81.79±3.52) and a corresponding large increase of the Type I error (FN: 18.22±3.75). At the same time, no significant changes for Type II errors, and the rate of true negative calls were observed (see Fig S1 in section SM-9). Overall, these results show that the LR_x_ statistics displays an excellent performance for the 2-person vs 2-person mixtures association test, and that it becomes more conservative when a common participant is looked for between 2-person and 3-person mixtures.

## 7. Matches and hits management

### 7.1. The match manager

Much work was devoted to the way results are presented to GENis users. In order to ease the analysis of query results for a given query profile (Q), GENis groups together target-matching profiles (T) by categories. For each one of these groups, hits can be listed by the mean fraction of shared alleles, the number of shared markers and the LR statistics defined to assess the statistical significance of the match (see Section UM-16 for implementation/operational details). Table 4 summarizes which statistics are considered by default for each kind of comparison.

**Table 4.** Analysis aim and default statistic employed for each query-target profile category combination (shorthand notation is used for LRs). Q: query profile, T: target profile, V: certain contributor profile (i.e. victim), U_i_: i-unknown contributor, X: profile of a common contributor, *n*_*Q*_ *(n*_*T*_): number of contributors of the query(target) profile.

| Query profile category | Target profile category | Default statistic | Looking for | Scenario testing |
| --- | --- | --- | --- | --- |
| S | S | $LR = \frac{1}{U_1}$ | identical twins<br>db consolidation | No |
| S | E<sub>1</sub> | $LR = \frac{Q}{U_1}$ | identification | No |
| S / E<sub>1</sub> | E | $LR = \frac{Q + U_1 + \dots + U_{n_T-1}}{U_1 + \dots + U_{n_T}}$ | contribution | Yes |
| | E<sub>v</sub> | $LR = \frac{Q + V + U_1 + \dots + U_{n_T-2}}{V + U_1 + \dots + U_{n_T-1}}$ | | |
| E | S / E<sub>1</sub> | $LR = \frac{T + U_1 + \dots + U_{n_Q-1}}{U_1 + \dots + U_{n_Q}}$ | | |
| E<sub>v</sub> | | $LR = \frac{T + V + U_1 + \dots + U_{n_Q-2}}{V + U_1 + \dots + U_{n_Q-1}}$ | | |
| E | E | $LR_x = \frac{X + U_1 + \dots + U_{n_T+n_Q-1}}{(U_1 + \dots + U_{n_T})(U_1 + \dots + U_{n_Q})}$ | common contributor | No |
| E<sub>v</sub> | | $LR_x = \frac{X + V_Q + U_1 + \dots + U_{n_T+n_Q-2}}{(U_1 + \dots + U_{n_T})(V_Q + U_1 + \dots + U_{n_Q-1})}$ | | |
| E<sub>v</sub> | E<sub>v</sub> | $LR_x = \frac{X + V_Q + V_T + U_1 + \dots + U_{n_T+n_Q-3}}{(V_T + U_1 + \dots + U_{n_T-1})(V_Q + U_1 + \dots + U_{n_Q-1})}$ | | |

### 7.2. The scenario-testing tool

Matching results can be prioritized by any of the above mentioned statistics. Noteworthy, GENis provides a *scenario testing framework*, inspired on the LRmix Studio software [13, 14] to further analyze reported matches in much detail. Different hypothesis and scenarios involving query (Q), target matching profiles (T) and *certain-group* profiles (C) matching the evidence sample at *moderate-level*, can be tested using this tool (see Section Manual UM-16.6 for usage examples).

### 7.3. Converting matches into hits

GENis provides a notification tray to inform each geneticist whether a match triggered by a third-party query involved any of his/her profiles (see Section UM-18 for further details on the notification system). In this way, when a potential hit is identified, the responsible geneticists are notified through the system. At this point the involved parties should contact each other to validate or refute the match. The *match manager module* will display the status of the reported match as: *pending* (the match should still be validated by one of the involved parties), *dismissed* (both parties agree on dismiss the match), *confirmed (*by both parties) or *in-conflict* (whenever one party confirmed the match, the other have dismissed it). See Section UM-16.4 for further operational details.

## 8. Multi-tier design

A GENis server can be locally installed to manage a single forensics lab’s DNA database. However, the main purpose of the GENis system is to be deployed in a multi-tier hierarchical architecture in order to share information between local, regional and/or national instances of the system.

The GENis tree-like network is built upon Laboratory and Registry type of nodes that are associated to participant forensics labs, where local profile databases are actually stored beingThese nodes the” leaves of the tree”. Regional Registries serve to integrate the information from affiliated laboratory nodes, and could typically be defined following judicial, geographical and/or administrative basis. In addition, a single master National Registry node can be deployed in order to coordinate and integrate data over the entire network.

Instance interconnectivity allows different nodes of the GENis ecosystem to communicate with each other through the hierarchical tree. Connectivity between instances uses SSL certificates to encrypt communications or can be configured to run over a VPN.

Uploading profiles information to a superior instance can be specified at profile’s entry/loading time. When a profile is sent to an upper instance it is stored in the corresponding category (mapping rules can be established to harmonize profile group and category definitions), and automatic queries are triggered in the higher instance. Whenever a match is detected, the information is transmitted to the involved lower instances in order to validate or discard the hit as explained en Section 7.3. Operational details of instance interconnectivity can be found at Section UM-21.

## 9. Discussion and conclusions

The use of DNA databases has had a profound impact on the ability to identify suspects linked to crime-scene evidence and to suggest/support investigation leads to relate evidence traces of unsolved cases.

In Argentina, following a request of the judiciary, the National Ministry of Science, Technology and Innovation undertook in 2014 an initiative to develop an open source software to assist in criminal investigations by identifying persons through biological evidence. Argentina is a federal country composed of 24 autonomous provinces that enact their respective laws. In the last 15 years, 19 provinces created their own genetic databases with different criteria regarding the types of crimes applicable and the defendant procedural status. With this in mind, GENis had a two-fold objective. First, it was aimed to provide a state-of-the-art tool for judicial institutions for storing and comparing DNA profiles in criminal cases. Secondly, GENis was intended to elicit comments and encourage debate in the experts’ community on the implementation of uniform protocols. Therefore, GENis could help to harmonize policies among provinces and facilitate data sharing strategies at the regional, national and international level.

GENis is a DNA information system that provides data integration capabilities at regional and/or national level expanding the scope, significance and capabilities of this kind of systems. the system-wide coordination of forensic information not only provides the possibility of running queries between otherwise independent local instances but also promotes the standardization of data structures and protocols.

As we have shown throughout the article, GENis is a highly flexible information system aimed to implement a very comprehensive DNA-related identification task. From a practical point of view, GENis integrates many ideas already developed by the international community, along with new statistical figures, into a single, unified and highly customizable framework that can accommodate many analysis workflows. Importantly, the system was entirely developed with strict adherence to open source policies to encourage the application of auditing procedures, and to warrant the integrity and ownership of the stored data. In addition, technical software specifications have been under exhaustive consistency examination by the Tools and Foundations for Software Engineering Lab [20], of the Computer Department of the University of Buenos Aires.

To date, GENis is already deployed in 18 different Argentinean provinces laboratories, and license agreements have been signed to expand it to new ones. A great effort was made in the training and assistance of technicians, geneticist and IT support teams. We have also made available a website (http://wiki.genis.sadosky.net) from where users can download useful information (handbooks, technical resources) and participate in forums to share operational feedback and suggest further developments.

## Acknowledgments

The initial requirement for GENis functionality map was set by the Argentine Society of Forensic Genetics (SAGF), the National Council of General Attorneys and the Federal Council of Criminal Policy. The program was developed by Fundacion Sadosky [21], a public-private institution in the orbit of the Argentine National Science and Technology Ministry (MINCyT), which coordinated an interdisciplinary team integrated by IT professionals, physicists, lawyers, molecular biologists and geneticists, representing very diverse institutional belongings. One remarkable asset of the project has been the synergy achieved through an active public-private cooperation along the whole development process. Researchers from various national universities (Buenos Aires, La Plata, Rosario, Córdoba, Tucumán, Quilmes, Comahue) have participated in different tasks, from the design of statistical models to the testing and validation of draft versions. Other R&D centers, like CONICET, Instituto Leloir Foundation and the Argentine Bioinformatics Society (A2B2C) have also taken part in the project bringing very useful contributions including training and technical guidance. For it’s part, the private sector was represented by a local IT company (Baufest) that was in charge of system design and coding. Additionally, the project benefitted from a continued support from the main national IT chambers (CESSI and CICOMRA).

Several Judicial institutions have also cooperated in the system development, including the Federal Court Board (JUFEJUS), the Laboratory of DNA Comparative Analysis of the Supreme Court of Justice of the Buenos Aires Province, the Forensic Genetic Service of the Supreme Court of Entre Rios Province, the Criminalistics and Forensic Sciences Research Institute within the scope the General Attorney’s Office of the Province of Buenos Aires, the Genetic Digital Fingerprint Service of the University of Buenos Aires and many others.

We appreciate the financial support from the MINCyT and Fundacion Sadosky. We would like to thank Mario Adaro (Institute of Technology of the Federal Court Board), Andrea Colussi (Forensic Medical Office of the National Supreme Court of Justice), Mercedes Lojo (DNA Comparative Analysis Laboratory of the Supreme Court of Justice of the Buenos Aires Province), Mariana Herrera, Walter Bozzo and Franco Marsico (National Bank of Genetic Data), Daniel Corach and Andrea Sala (Genetic Digital Fingerprint Service of the University of Buenos Aires), Cesar Guida and Elina Francisco (Criminal Investigation and Forensic Sciences Institute of the General Attorney’s Office of Buenos Aires province), Cecilia Miozzo (NOA Regional Forensic Genetic Laboratory of the Supreme Court of Justice of Jujuy province), Alejandra Guinudinik (Forensic Molecular Biology Service of the General Attorney’s Office of Salta province), Pedro Villagran (Forensic Investigations Center of the General Attorney’s Office of Salta province), Silvia Vanelli Rey (North Patagonia Regional Forensic Genetic Laboratory of the General Attorney’s Office of Rio Negro province), Cecilia Bobillo (Forensic Genetic Laboratory of the General Attorney’s Office of La Pampa province), María Beatriz Vazquez (Forensic Investigations Laboratory of the Supreme Court of San Juan province), Haydee Fariña (Forensic Medical Service of the Supreme Court of Neuquen province), Gabriela Lamparelli (Forensic Sciences Institute of the Supreme Court of Chaco province), Juan José Belzki (Forensic Investigation Center of the Supreme Court of Formosa province), Agustina Dorigón Lezana (Forensic Genetic Cabinet of the Supreme Court of Santiago del Estero province), Victor Moloeznik (General Attorney’s Office of Santa Fe province), María Consuelo Martí (Chemical Laboratory of the General Attorney’s Office of Santa Fe province), Juan José Galvez (Forensic Medical Institute of the Supreme Court of Corrientes province), Diego Rinaldi (General Attorney’s Office of Corrientes province), Noelia Massari (Regional Forensic Investigations Laboratory of the General Attorney’s Office of Chubut province), Miguel Rubio (Forensic Genetic Cabinet of the Supreme Court of Tucuman province), Nestor Basso and Lidia Vidal Rioja (CONICET), Hortensia Cano (South Patagonia Regional Forensic Genetic Laboratory of the Supreme Court of Justice of Santa Cruz province), Inés Aparici and Jessica Name (Supreme Court of Tierra del Fuego Province), Cristina Buslje (Fundación Instituto Leloir, A2B2C), Sebastian Uchitel, Hernán Melgratti and Víctor Braberman (Buenos Aires University, CONICET), Gustavo Parisi (National Quilmes University), Paula Venosa (La Plata University), Sandra Furfuro (Cuyo University), Manuel Aybar (Tucumán University), Carlos de la Vega (Cordoba University), Juan Luzuriaga and Agustina Buccella (Comahue University) and many others for fruitful discussions.

## Footnotes

1 Fundación Sadosky holds all rights over GENis software and authorizes the publication of this article.

2 European Network of Forensic Sciences Institutes

3 International Society of Forensic Genetics

4 Operational arrangements have already been addressed in Argentina with the national blockchain authority (BFA-Blockchain Federal Argentina) to activate this option upon request of the court system.

5 Data from http://www.ffyb.uba.ar/shdg/batabase-2009?en,,mnu-e-46-7-mnu- Dr D.Corach Private communication.

