## Supplementary Material for "GENis, an open-source multi-tier forensic DNA information system"

### 1. Genotype probability models

In this section we summarize the main ideas of the assumed genotype probability models. We refer to [Fung2008] for a more in-depth discussion regarding this topic.

For the general case of a population that comprises various subpopulations, if evolutionary equilibrium is assumed, the allele proportions, $\{p_i\}$, can be modeled considering a Dirichlet distribution [Wright1951].

$$P(\{p_i\}) = \frac{\Gamma(\gamma)}{\prod_i \Gamma(\gamma_i)} \prod_i p_i^{\gamma_i} \qquad [1]$$

where $\Gamma(\gamma)$is a gamma function, $p_i$ is the proportion of allele $A_i$ and

$$\gamma_i = \frac{(1-\theta)p_i}{\theta} \quad , \quad \gamma = \sum_i \gamma_i = \frac{(1-\theta)}{\theta} \qquad [2]$$

The parameter $0 < \theta < 1$ is called the *inbreeding coefficient.* It is a positive number that can be regarded as a measure of the variation in subpopulation allele proportions (i.e. the larger $\theta$, the more *flat* the probability distribution will be)

As stated in [Fung2008], the probability of seeing allele $A_i$ occurring $m_i$ times in a set of $m$ alleles, given the proportions $p$'s, will follow a multinomial distribution:

$$P\left(\prod_i A_i^{m_i} \middle| \{p_i\}\right) = \frac{(\sum_i m_i)!}{\prod_i (m_i)!} \prod_i p_i^{m_i} \qquad [3]$$

Then, it follows the marginal probability can be written as

$$P\left(\prod_i A_i^{m_i}\right) = \frac{\Gamma(\gamma)}{\Gamma(\gamma+m)} \prod_i \frac{\Gamma(\gamma_i + m_i)}{\Gamma(\gamma_i)} \qquad [4]$$

Importantly, Equation (4) can be used to assess the probability of seeing allele $A_i$ occurring $m_i$ times in a set of $m$ alleles. In particular, when applied to a simple genotype $(A_i, A_j)$ Eq (4) implies that:

$$P(A_i, A_j) \equiv \begin{cases} p_i^2 + p_i(1-p_i)\theta & i = j \\ 2p_ip_j(1-\theta) & i \neq j \end{cases} \quad [5]$$

The NRCII Recommendation 4.1 adopts a somehow more conservative estimation of the genotype probability for the herterozygote case (compatible with the Hardy-Wainberg equilibrium hypothesis).

NRCII 4.1

$$P(A_i, A_j) \equiv \begin{cases} p_i^2 + p_i(1-p_i)\theta & i = j \\ 2p_ip_j & i \neq j \end{cases} \quad [6]$$

At this point it is worth remembering that joint and conditional probabilities are related by this basic equation:

$$P(X, Y) = P(X|Y)P(Y)$$

Hence, it is easy to show that the multivariate probability function associated to a multi-set of alleles described in Equation [4] implies that

$$P(A_i | y\ A_i \text{ alleles among } n \text{ alleles}) = \frac{y\theta + (1-\theta)p_i}{1 + (n-1)\theta} \quad [7]$$

Eq [7] can be used to estimate the probability of observing an allele multi-set **given** the observation of another one. This is a handy formula, as it allows us to quantify the probability of observing a random person, *U*, from the population of interest having the same genotype than the already observed one for suspect *S*: $P(U = A_iA_j | S = A_iA_j)$

This conditional probability form is the one considered in the NRCII recommendation 4.10 to estimate genotype probabilities from allelic ones:

NRCII 4.10

$$P(A_i, A_j) \equiv \begin{cases} \frac{[2\theta + (1-\theta)p_i][3\theta + (1-\theta)p_i]}{(1+\theta)(1+2\theta)} & i = j \\ \frac{2[\theta + (1-\theta)p_i][\theta + (1-\theta)p_j]}{(1+\theta)(1+2\theta)} & i \neq j \end{cases} \quad [8]$$

### 2. New allele frequencies

GENis provides 4 alternatives to set frequencies attributed to new allele values:

$$f_{min} = \begin{cases} f_0 \text{ (fixed value)} \\ \frac{5}{2N} \text{ (NRC II)} \\ 1-\alpha^{1/2N} \quad \text{[Weir1992]} \\ 1-\left[1-(1-\alpha)^{\frac{1}{C}}\right]^{\frac{1}{2N}} \quad \text{[Budowle1996]} \end{cases}$$

The first option considers a user provided fixed value, $f_0$ . The second one is compliant with NRCII recommendation and takes into account the allelic database size (N is the number of sampled people). The last two options provide two alternative assessments of the minimum allele frequency for a database, estimated with $100(1-\alpha)$ confidence. *C* is the number of common alleles which can be estimated from the level of heterozygosity of the characterized marker in the population [Budowle1996].

#### 3. Binary model for DNA profiles

GENis uses a binary approximation to model DNA profiles, produced through the Polymerase Chain Reaction amplification of regions of the genome that contain Short Tandem Repeats (STRs). The following table summarizes the necessary definitions of our model.

| Concept | Data type | Description | Definition | Example |
|---|---|---|---|---|
| **Locus, marker or system** | string | genomic position of a given STR used for identification | | D3S1358, TPOX,… |
| **Allele** | numeric | Number of STR repetitions | $x \equiv \frac{n}{100}, n \geq 100 \in \mathbb{N}$ | 13, 9.3 |
| | | Obligated allele for matching purposes (modifier) | [*x*] | [6] |
| **Genotype** | Tuple | | Locus->(Allele, Allele,…) | D3S1358 → (28.5, 22). |
| **Genotypifica-tion** | List | List of genotypes (no duplicated loci) | G={Locus->(Allele, Allele,…), Locus->(Allele, Allele,…),…} | {D3S1358 → (28.5, 22), D16S539→(5), Penta E→(16.3,18),…} |
| **Profile** | Structure | A DNA profile is a genotypification identified by a unique id | Id + Genotypification | [1678-DH, {D3S1358 → (28.5, 22), D16S539→(5),…}] |

Definition of GENis binary model

### 4. Bayesian framework

#### 4.1. Likelihood calculations

GENis allows comparing conditional probabilities under alternative scenarios in order to quantitatively examine different hypothesis regarding the source of crime scene samples. For each analyzed system likelihood ratios are estimated as

$$LR = \frac{P(ev\ |H_1)}{P(ev\ |H_0)} \qquad [9]$$

The evidence, *ev,* is made up of the allele multi-set of available genotyped profiles. $H_1$ and $H_0$ are the two competing hypothesis that propose different participation patterns of genotyped ($S_1$,…, $S_n$)and unknown ($U_1$,…, $U_m$)contributors to a given evidence profile *M*.

Following [Curran2005] and [Haned2012], the general form of the evidence probability under $H_i$can be written as

$$P(ev\ |H_i) = \sum_j P\left(M, K, V, U_j \middle| H_i\right) \qquad [10]$$

where *M* represents the crime-scene sample allele set, *K* is the allele multi-set of genotyped persons that, according to $H_i$ contributed to the sample. *V* is the allele multi-set of genotyped persons that, according to $H_i$ did not contribute to the sample. $U_j$ represents the j-putative allele multi-set of the $n_u$ unknown contributors, proposed by $H_i$.

Using basic probability considerations, each term of equation [10] can be written as

$$\begin{aligned} P\left(M, K, V, U_j \middle| H_i\right) &= P\left(M \middle| K, V, U_j, H_i\right) P\left(K, V, U_j \middle| H_i\right) \\ &= P\left(M \middle| K, U_j, H_i\right) P\left(K, V, U_j \middle| H_i\right) \end{aligned} \qquad [11]$$

Therefore,

$$P(ev\ |H_i) = \sum_j P\left(M \middle| K, U_j, H_i\right) P\left(K, V, U_j \middle| H_i\right) \qquad [12]$$

where the sum iterates over the different sets of $n_u$ unknown individuals that could have contributed to sample *M.* In order to generate the possible $\{U_j\}$ sets, GENis considers all possible combination with repetition of 2 $n_u$ alleles, taken from the available *k* allele values reported for the locus of interest.

The genotype probability $P(K, V, U_j | H_i)$ can be straightforwardly estimated considering Eq[7]. On the other hand, the probability $P(M | K, U_j, H_i)$ of observing sample *M*, assuming a given contributors scenario, can be obtained taking into account *drop-out* (allele losses) and *drop-in* (allele contamination) probability values [Curran2005, Haned2012]. Let $\delta$ be the allele multi-set of the sample contributors (*K* and $U_j$) that are absent in *M* (i.e. alleles affected by drop-out events). Let $\chi$ be the allele multi-set present in sample *M*, but not included in any contributor profile (i.e. contamination alleles). Finally, let $\rho$ represents alleles in *K* and $U_j$ that are also found in *M.* The desired probability can be estimated as:

$$P(M | K, U_j, H_i) = p_\delta p_\rho p_\chi \qquad [13]$$

with

$$p_\delta = \prod_{A_i \epsilon \delta} (p_{out})^{n_{A_i}} \qquad p_\rho = \prod_{A_i \epsilon \rho} (1 - {p_{out}}^{n_{A_i}}) \qquad p_\chi = \begin{cases} 1 - p_{in} \ \ si \ |\chi| = 0 \\ p_{in}^{|\chi|} \prod_{A_i \epsilon \chi} {p_{A_i}}^{n_{A_i}} \end{cases}$$

$p_{out}$ and $p_{in}$ are the dropout and dropin probability values respectively. $n_{A_i}$ and $p_{A_i}$ are the multiplicity and the population frequency of allele $A_i$ respectively.

### 5. Automatic estimation for the number of contributors

In order to provide an estimator for the number of contributors ($n^*$) of a given sample GENis follows the strategy proposed in [Haned2011] and maximizes the probability that the mixture *M* came from *n* random contributors, as a function of *n:*

$$n^* = \underset{1 \le n \le 5}{\operatorname{argmax}} \ P(M, U_1, \ldots, U_n) \qquad [14]$$

with

$$P(M, U_1, \ldots U_n) = \sum_j P\left(M \middle| U_1^{(j)}, \ldots, U_n^{(j)}\right) P(U_1^{(j)}, \ldots, U_n^{(j)}) \qquad [15]$$

The index *j* iterates over possible sets of *x* unknown contributors and $U_i^{(j)}$ is the genotype of the i-random man contributors of the j-set.

### 6. Matching stringency rules

GENis adopted the three stringency level scheme suggested in Section 5.4.2 of the ENFSI Best practice manual for DNA pattern recognition and comparison document. The following table summarizes the stringency criteria that can be used to define matching rules for DNA profiles

| Stringency | Description | Locus comparison scheme |
| --- | --- | --- |
| **High** | all alleles of **every locus** present in one DNA profile must also be present in the matching DNA profile | 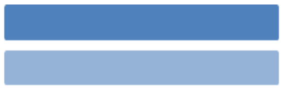 |
| **Moderate** | of two DNA profiles, the alleles of a locus with the least number of alleles must be present in the corresponding locus of the other DNA profile | 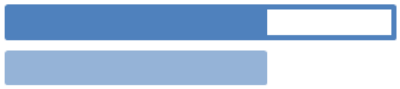 |
| **Low** | in each locus compared between two DNA profiles, at least one allele of that locus must be present in the other DNA profile | 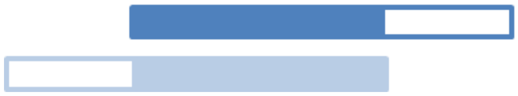 |

### 7. Common contributor likelihood

Let's assume that, for a given locus *s*, a pair of samples, $M_1$ and $M_2$ (having $n_1 \geq 2$ and $n_2 \geq 2$ contributors respectively) match at low stringency levels. We wonder if it is safe to assume that a common contributor exists. GENis considers all the pair combinations (with repetitions) of shared alleles to build *X,* the set of putative common contributors.

We start by considering the probability of finding the available evidence under the hypothesis $H_x$ that sustains the existence of a common unknown contributor, member of $X = \{X_1,..X_{nx}\}$, in both mixtures:

$$P(ev|H_x) = \sum_{j,k,l} P(M_1, M_2, K_1, K_2, X_j, Y_{1k}, Y_{2l}) \quad [16]$$

$K_i$ is the allele multi set of the $n_{ki}$ known contributors of mixture *i* ($i$=1,2), $X_j$ represents the alleles of the putative *j*-unknown common contributor, and $Y_{ik}$ is the *k*-allele multi set for the $n_i$ - $n_{ik}$ – 1 putative unknown contributors that, along with $X_j$, appear in sample *i*. As we show in SM-4.2, Equation (1) can be further casted into the following expression:

$$P(ev|H_x) = \sum_{j} P_j(ev|H_x) \quad [17]$$

where

$$P_j(ev|H_x) \equiv P(K_1, K_2, X_j) \sum_{k,l} P(M_1|K_1, X_j, Y_{1k}) P(M_2|K_2, X_j, Y_{2l}) P(Y_{1k}Y_{2l}|K_1, K_2, X_j)$$

On the other hand, the likelihood supporting hypothesis $H_0$, that both samples were in fact statistically independent, can be written as:

$$P(ev|H_0) = \left(\sum_{l} P(M_1, K_1, U_{1l})\right)\left(\sum_{l} P(M_2, K_2, U_{2l})\right) \quad [18]$$

where $U_{il}$ is the *l*-allele multi set of the $n_i$ - $n_{ki}$ unknown contributors of sample-i under $H_0$

In order to statistically assess whether a common contributor had participated in both mixture samples Eqs (2) and (3) can be combined into the following likelihood ratio:

$$LR^{(s)} = \frac{P(ev|H_x)}{P(ev|H_0)} = \sum_{j=1}^{n_x} LR_j^{(s)} \quad [19]$$

where $LR_j$ is the contribution to the likelihood ratio of the putative common contributor $X_j$:

$$LR_j^{(s)} = \frac{P_j(ev|H_x)}{\sum_k P(M_1, K_1, U_{1k}) \sum_l P(M_2, K_2, U_{2l})} \quad [20]$$

Multivariate probabilities and likelihood factors appearing in the last equations can be estimated as detailed in supplementary material SM-4.1. Noticeably, Eq [5] provides an estimation of the impact of each putative common contributor to the overall LR value. In this way, $LR_j^{(s)}$ could be used to rank them in terms of their relative relevance. Moreover, the $\max_j(LR_j^{(s)})$ value can be used as an alternative proxy of the sought sample-pair association index.

Finally, assuming statistical independence between markers, the overall LR statistics results:

$$LR_x = \prod_s LR^{(s)} \qquad [21]$$

### 8. Classification performance

The classification performance induced by the $LR_x$ statistics can be quantified considering two figures of merit: the F1 score and the Youden's J statistics [9]. The former is the harmonic mean of the precision and sensitivity of the classification system. The second one aims to capture the performance of dichotomous classification heuristics taking into account the specificity and the sensitivity of the test:

$$F_1 = 2\,\frac{precision.recall}{precision + recall} = \frac{2\,TP}{2\,TP + FP + FN} \qquad [22]$$

$$J = sensitivity + specificity - 1 = \frac{TP}{TP + FN} + \frac{TN}{TN + FP} - 1 \qquad [23]$$

TP, FP, TN, and FN stand for: true positive, false positive, true negative, and false negative values respectively.

### 9. **Supplementary Figures**

Figure S1 Upper panel: Density distribution functions for the $LR_x$ statistics are shown for 2-contributor vs 3-contributor's mixtures association tests for case (black lines) and control (gray lines) independent pairs.. The respective confusion matrix is displayed as an inset. Lower panel: Classification performance of the $LR_x$ statistics to discriminate case and control profile pairs, as a function of the number of considered markers. F1 and Youden's J statistics are shown using solid and empty black symbols respectively. The fraction of null $LR_x$ obtained for independent control profile pairs is shown using red empty squared symbols and a dashed red line.

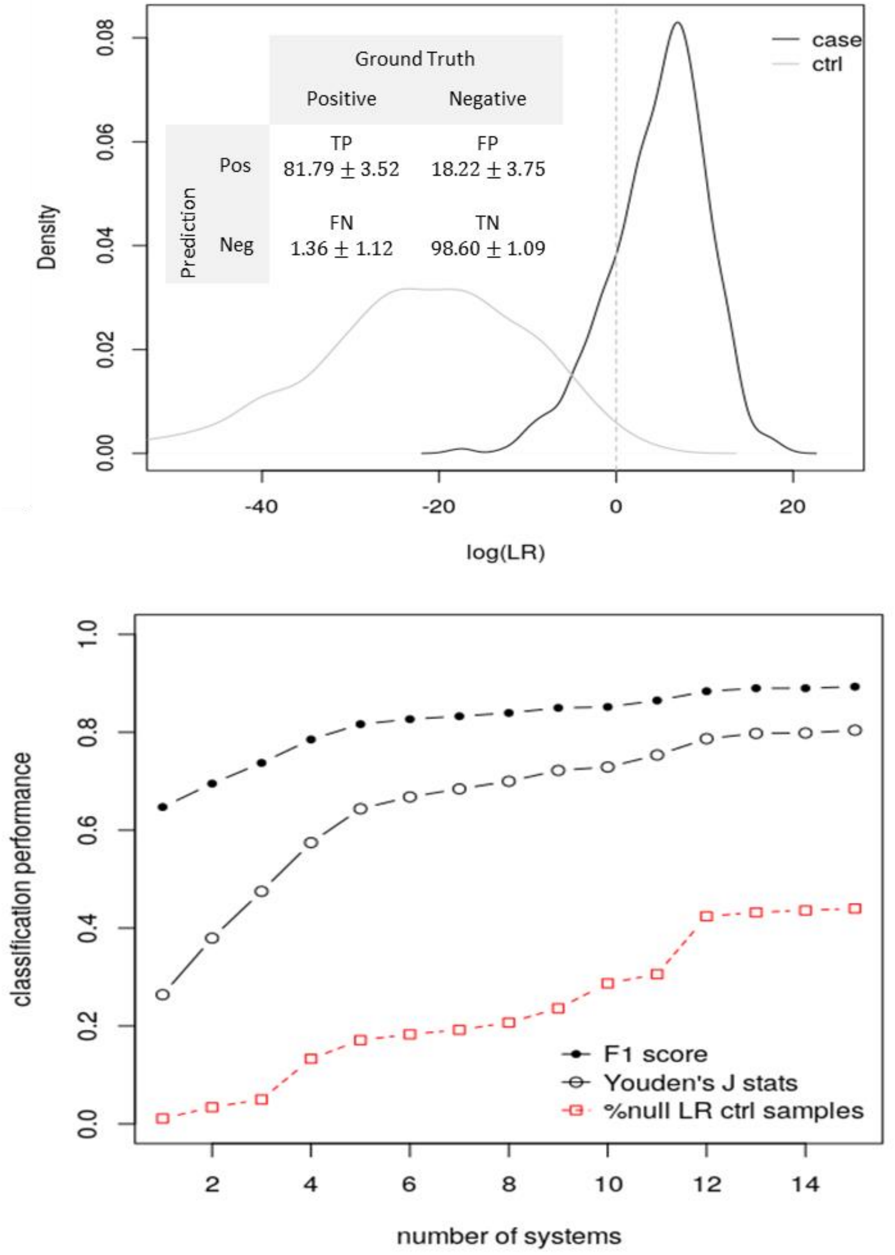

[24]

**Table SM-T1.** Permissions that can be granted to different roles and associated operations are detailed.

| Permission | Allowed operations |
|---|---|
| Category management | Registration of category groups.<br>Modification of category groups.<br>Reading category descriptions.<br>Reading of groups and categories.<br>Unsubscribe categories.<br>Modification of categories.<br>Subscribe categories.<br>Unsubscribe category groups. |
| Profile management | Reading of groups and categories.<br>Association of genetic profiles.<br>Subscribe genetic profiles.<br>Subscribe genetic analysis.<br>Obtaining a token to upload images.<br>Notification reading.<br>Reading genetic profiles.<br>Association of genetic profiles.<br>Upload of images.<br>Subscribe electropherograms.<br>Reading the sample data.<br>Reading STR Kits. |
| Reading of allele frequency tables | Reading of allele frequency tables.<br>Subscription of allele frequency tables.<br>Modification of allele frequency tables.<br>Notification reading. |

| | |
|---|---|
| User administration | User reading.<br>Modification of users.<br>Modification of user status. |
| Batch load | Profile batch upload.<br>Search for profiles coming from batch load.<br>Acceptance and rejection of profiles coming from batch load.<br>Acceptance and rejection of profiles coming from batch load.<br>High profile metadata from batch load.<br>Reading and search of profiles coming from batch load. |
| Profile search & Scenario management | Reading scenarios.<br>Subscribe scenarios.<br>Unsubscribe scenarios.<br>Reading the sample data.<br>Validation of scenarios.<br>Modification of scenarios.<br>Reading genetic profiles.<br>Match reading.<br>Laboratory reading. |
| Laboratory/geneticists administration | Modification of laboratories.<br>Laboratory registration.<br>Countries reading.<br>Provinces reading.<br>Laboratory reading.<br>Reading of geneticists.<br>Modification of geneticists.<br>Registration of geneticists.<br>Notification reading. |
| Profile data management | Subscription of the sample data. |

| | |
|---|---|
| | Reading category descriptions.<br>Reading of geneticists.<br>Reading of groups and categories.<br>Modification of the sample data.<br>Notification reading.<br>Reading types of biological material.<br>Laboratory reading.<br>Obtaining a token to upload images.<br>Unsubscribe profiles.<br>Upload of images.<br>Reading the sample data.<br>Reading types of crimes. |
| Types of biological material | Reading types of biological material.<br>High of types of biological material.<br>Modification of types of biological material.<br>Lower types of biological material. |

**Table SM T1** Marker kits readily available in GENis

| Kits | STR markers |
|---|---|
| ENFSI/ EuropeanUnion | D10S1248,D12S391, D18S51,D1S1656, D21S11,D22S1045. D2S441,D3S1358, D8S1179,FGA, THO1,vWA |
| Globalfiler | SE33, Amelogenin, CFS1PO, D10S1248, D12S391, D13S317, D16S539, D18S51, D19S433, D1S1656, D21S11, D22S1045, D2S1338, D2S441, D3S1358, D5S818, D7S820, D8S1179, DYS391, FGA, THO1, TPOX, vWA, YIndel |
| Identifiler | D8S1179, D21S11, D7S820, CSF1PO, D3S1358, THO1, D13S317, D16S539, D2S1338, D19S433, vWA, TPOX, D18S51, Amelogenin, D5S818, FGA |
| Interpol | SE33, Amelogenin, CSF1PO, D10S1248, D12S391, D13S317, D16S539, D18S51, D19S433, D1S1656, D21S11, D22S1045, D2S1338, D2S441, D3S1358, D5S818, D7S820, D8S1179, FGA, PentaD, PentaE, THO1, TPOX, vWA |
| Investigator Decaplex SE Kit | SE33,Amelogenin, D16S539,D18S51, D19S433,D21S11, D2S1338,D3S1358, D8S1179,FGA,THO1, vWA |
| Investigator ESSplex Plus Kit | Amelogenin, D10S1248, D12S391, D16S539, D18S51, D19S433, D15S1656, D21S11, D22S1045, D2S1338, D2S441, D3S1358, D8S1179, FGA, THO1, vWA |
| Investigator ESSplex SE Plus Kit | SE33, Amelogenin, D10S1248, D12S391, D16S539, D18S51, D19S433, D1S1656, D21S11, D22S1045, D2S1338, D2S441, D3S1358, D8S1179, FGA, THO1, vWA |
| Investigator HDplex Kit | SE33,Amelogenin, D10S2325,D12S391, D18S51,D21S2055, D2S1360,D3S1744, D4S2366,D5S2500, D6S474,D7S1517, D8S1132 |
| Investigator Hexaplex ESS Kit | Amelogenin,D10S1248, D12S391,D1S1656, D22S1045, D2S441, THO1 |
| Investigator IDPlex Plus Kit | Amelogenin, CSF1PO, D13S317, D16S539, D18S51, D19S433 D21S11, D2S1338, D3S1358, D5S818, D7S820, D8S1179, FGA, THO1, TPOX, vWA |
| Investigator Nonaplex ESS Kit | SE33, Amelogenin, D10S1248, D12S391, D18S51, D1S1656, D21S11, D22S1045, D2S441, D3S1358, D8S1179, FGA, THO1, vWA |
| Investigator Triplex AFS | FGA, SE33, Amelogenin |

| | |
|---|---|
| QS Kit | |
| Investigator Triplex<br>DSF Kit | D3S1358, FGA, SE33 |
| Minifiler | D13S317,D7S820, Amelogenin,D2S1338, D21S11,D16S539, D18S51, CSF1PO, FGA |
| NGM | Amelogenin,D10S1248, D12S391,D16S539, D18S51,D19S433, D1S1656,D21S11, D22S1045,D2S1338, D2S441,D3S1358, D8S1179,FGA,THO1, vWA |
| NGMSelect | SE33, Amelogenin, D10S1248, D12S391, D16S539, D18S51, D19S433, D1S1656, D21S11, D22S1045, D2S1338, D2S441, D3S1358, D8S1179, FGA, THO1, vWA |
| Poweplex 16 | D3S1358, THO1, D21S11, D18S51, PentaE, D5S818, D13S317,D7S820, D16S539, CSF1PO, PentaD, Amelogenin,vWA,D8S1179, TPOX, FGA |
| Poweplex 18D | D2S1338, TPOX, FGA, D8S1179,vWA, Amelogenin,D3S1358, THO1,PentaE,D18S51, D21S11, D5S818,CSF1PO, D7S820,D13S317,D16S539, PentaD, D19S433 |
| Powerplex 2.1 | D18S51, D21S11, D3S1358, D8S1179, FGA, PentaE, THO1, TPOX, vWA |
| Powerplex 21 | Amelogenin, D3S1358, D1S1656, D6S1043, D13S317, PentaE, D16S539, D18S51, D2S1338, CSF1PO, PentaD, THO1, vWA, D21S11, D7S820, D5S818, TPOX, D8S1179, D12S391, D19S433, FGA |
| Powerplex 1.2 | CSF1PO, D13S317, D16S539, D5S818, D7S820, THO1, TPOX, vWA |
| Powerplex CS7 | F13B, F13A, FPS, LPL, PentaC, PentaD, PentaE |
| Powerplex ES | SE33, Amelogenin, D18S51, D21S11, D3S1358, D8S1179, FGA, THO1, vWA |
| Powerplex ESI 16 (Fast) | Amelogenin, D10S1248, D12S391, D16S539, D18S51, D19S433, D1S1656, D21S11, D22S1045, D2S1338, D2S441, D3S1358, D8S1179, FGA, THO1, vWA |
| Powerplex ESI 17 (Fast) | SE33,Amelogenin, D10S1248,D12S391, D16S539,D18S51, D19S433,D1S1656, D21S11,D22S1045, D2S1338,D2S441, D3S1358, D8S1179, FGA, THO1, vWA |

| | |
|---|---|
| Powerplex ESX 16 (Fast) | Amelogenin, D10S1248, D12S391, D16S539, D18S51, D19S433, D1S1656, D21S11, D22S1045, D2S1338, D2S441, D3S1358, D8S1179, FGA, THO1, vWA |
| Powerplex ESX 17 (Fast) | Amelogenin,D10S1248, D12S391,D16S539, D18S51,D19S433, D1S1656,D21S11, D22S1045,D2S1338, D2S441,D3S1358, D8S1179,FGA,THO1, vWA, SE33 |
| Powerplex Fusion | Amelogenin, CSF1PO, D10S1248, D12S539, D13S317, D16S539, D18S51, D19S433, D1S1656, D21S11, D22S1045, D2S1338, D7S820, D8S1179, DYS391, FGA, PentaD, PentaE, THO1, TPOX, vWA |
| Powerplex S5 | FGA, D8S1179, THO1, D18S51, Amelogenin |
| Profiler | Amelogenin, CSF1PO, D13S317, D3S1358, D5S818, D7S820, FGA, THO1, TPOX, vWA |
| Profiler + | Amelogenin, D13S317, D18S51, D21S11, D3S1358, D5S818, D7S820, D8S1179, FGA, vWA |
| SE-filer | SE33, Amelogenin, D16S539, D18S51, D19S433, D21S11, D2S1338, D3S1358, D8S1179, FGA, THO1, vWA |
| SGM + | Amelogenin, D16S539, D18S51, D19S433, D21S11, D2S1338, D3S1358, D8S1179, FGA, THO1, vWA |
| Sinofiler | CSF1PO,D12S391, D13S317,D16S539, D18S51, D19S433,D21S11, D2S1338,D3S1358, D5S818, D6S1043,D7S820, D8S1179, FGA, vWA |
| SoleKit | CSF1PO, TPOX, FGA, THO1 |
| USA (CODIS core-loci) | Amelogenin,CSF1PO, D13S317,D16S539, D18S51, D21S11, D3S1358, D5S818, D7S820, D8S1179, FGA, THO1, TPOX, vWA |
| Verifiler | D10S1248,D12S391, D19S433,D1S1656, D22S1045,D2S1338, D2S441, D6S1043, THO1 |
